# Forest demography and biomass accumulation rates are associated with transient mean tree size vs density scaling relations

**DOI:** 10.1101/2023.12.11.571032

**Authors:** Kailiang Yu, Han Y.H. Chen, Arthur Gessler, Thomas A.M. Pugh, Eric B. Searle, Robert B. Allen, Hans Pretzsch, Philippe Ciais, Oliver L. Phillips, Roel J.W. Brienen, Chengjin Chu, Shubin Xie, Ashley P. Ballantyne

## Abstract

Linking individual and stand-level dynamics during forest development reveals a scaling relationship between mean tree size and tree density in forest stands, which integrates forest structure and function. However, the nature of this so-called scaling law and its variation across broad spatial scales remains unquantified and its linkage with forest demographic processes and carbon dynamics remains elusive. Here we develop a theoretical framework and compile a broad-scale dataset of long-term sample forest stands (n = 1433) from largely undisturbed forests to examine the association of temporal mean tree size vs density scaling trajectories (slopes) with biomass accumulation rates and the sensitivity of scaling slopes to environmental and demographic drivers. The results empirically demonstrate a large variation of scaling slopes, ranging from -4 to -0.2, across forest stands in tropical, temperate and boreal forest biomes. Steeper scaling slopes are associated with higher rates of biomass accumulation, resulting from a lower offset of forest growth by biomass loss from mortality. In North America, scaling slopes are positively correlated with forest stand age and rainfall seasonality, thus suggesting a higher rate of biomass accumulation in younger forests with lower rainfall seasonality. These results demonstrate the strong association of the transient mean tree size vs density scaling trajectories with forest demography and biomass accumulation rates, thus highlighting the promise of leveraging forest structure properties to predict forest demography, carbon fluxes and dynamics at broad spatial scales.

**Significance Statement:** Mean tree size vs density scaling relationships are thought to predict forest function at broad spatial scales. Here we develop a theoretical framework based upon demographic processes and empirical evidence from forest inventory data to demonstrate a strong association of the transient mean tree size and density scaling trajectories (slopes) with forest demography and biomass accumulation rates. This strong association is pervasive across forest biomes and suggests a negative relationship between scaling slope and biomass accumulation rate (resource availability). Our results highlight the promise of leveraging forest structure (i.e., inferred from high resolution remote sensing data or fused into size-structured demographic models) to evaluate forest demography, carbon fluxes and dynamics at broad spatial scales.

## Introduction

A central research focus in forest ecology has been examining the spatiotemporal dynamics of forest functions that affect carbon stocks and fluxes (1–3). The linkage of forest structure and function is thought to enable an improved understanding or prediction of the demography and carbon dynamics for the entire forest by inferring its structural properties (i.e., size and/or density) (4–6). Discerning this linkage between forest structure and carbon dynamics holds promise for understanding forest carbon sinks at broad or global scale by leveraging off emerging advanced tools such as high resolution satellite remote sensing data (7, 8) and size-structured forest demographic models (6, 9–11). However, a theoretical framework with empirical evidence linking forest structure and function is still lacking, especially at broad spatial scales.

Mean tree size and density scaling relations provide a link between forest structure and function. The underlying rationale is based on the demographic process that in dense stands increased size of some individual trees comes at the expense of other individuals due to competition for finite resources (e.g. space, light, water, nutrients), thus leading to a decrease in tree density and an increase in mean tree size during forest development (12, 13). These transient dynamics during the life cycle of a forest can be described by a scaling law between mean individual tree size (𝑀̅, aboveground biomass, hereafter referred to as tree size) and the density (𝑁) of living trees in a community (𝑀̅ = 𝑘 𝑁*^α^*) at a given forest stand age, where 𝑘 is the coefficient and *α* is the scaling exponent (4, 6, 14). At macroecological scales some studies suggest an Euclidean scaling exponent of -3/2 (14, 15), but others highlight the possibility of a fractal scaling exponent of -4/3 (16, 17) (Fig. 1a). Further studies at local scales have found substantial variation of scaling exponents, which appear to be subject to environmental and biological constraints, or argued for these based on theory (18, 19). However, these studies adopted a spatial approach to determine *α*, using data of mean tree size and density from many stands spread across a large area. This spatial approach may hide both variations in scaling trajectories and key factors responsible for this variation if different forest types or locations are used in space for time analyses (4, 15). Moreover, by not directly isolating temporal variations in forest structure, spatial approaches do not provide insight into how this mean size vs density scaling trajectory may affect forest demography and carbon sequestration at forest stand scales under changing environmental conditions (5, 15, 20, 21). To untangle these factors a new approach calculating *α* based on temporal changes in stand structure is needed.

**Fig. 1.**
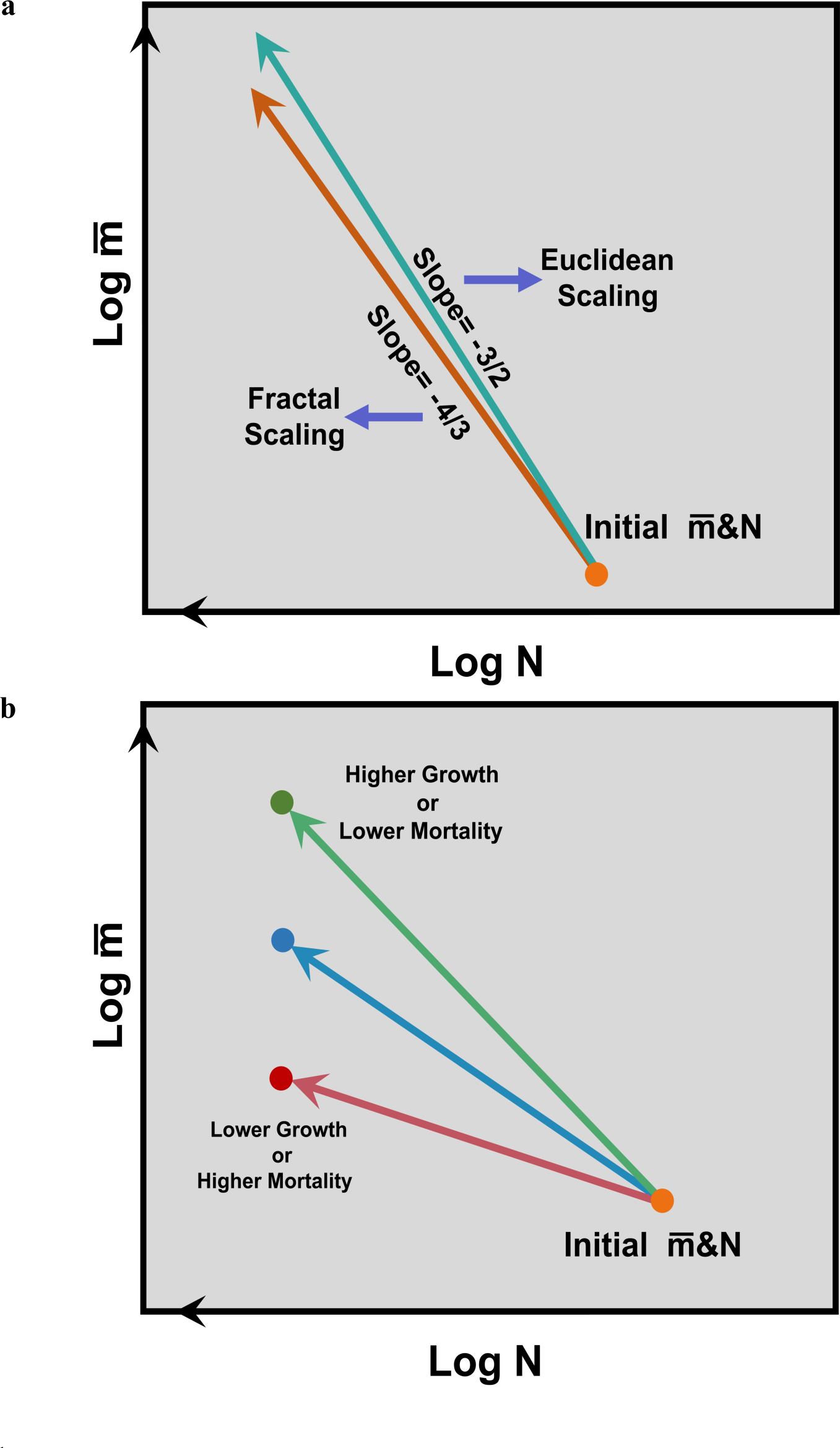
A conceptual framework on the transient mean tree size vs density scaling law: the scaling trajectory and its association with forest demography and biomass accumulation rates. **(a)** The mean tree size (aboveground biomass) vs density scaling law is initially proposed and applied in even sized community whereby trees compete for the constant space and/or resources. At macroecology scales (i.e., by using large scale and spatial data of mean tree size and density) the scaling law predicts a scaling exponent (𝛼) of -3/2 (Euclidean scaling) or -4/3 (fractal scaling), whereby the isometric model considers plants as 3-dimensional solids (Euclidean scaling) and the allometric model considers the geometric structure of plant as a fractal. As such, this spatial approach does not allow to link the demography with 𝛼 at forest stand or local scales. **(b)** The transient mean tree size vs density scaling trajectory or temporal 𝛼 is expected to be linked with forest demography – growth and mortality at forest stand or local scales. Here the transient mean tree size vs density scaling trajectory is temporally fitted using temporal series of data on mean tree size and density within each forest stand. As conceptualized based on demographic processes, when the mortality rate remains constant, the higher forest growth would lead to higher mean size and thus steeper mean tree size vs density scaling trajectory with more negative value of scaling exponent and higher biomass accumulation rate. By comparison, the lower growth rate would lead to less negative value of scaling exponent. Similarly, in the scenario of assuming the constant forest growth, the higher mortality rate would decrease the time to reach the decreased tree density (i.e., 200 ind ha^1-^), thus leading to the lower mean size, a less negative value of scaling exponent and the lower biomass accumulation rate. By comparison, the lower mortality rate would lead to more negative value of scaling exponent. See the conceptual model for mathematic derivations in details in Methods.

Here we propose a conceptual model based on ecological theory (see Methods) to illustrate our hypotheses that the temporal transient mean tree size vs density scaling exponent (slope, *α*) (12, 22) is related to the rate of biomass change over time and that growth and biomass loss from mortality jointly (or growth and biomass loss ratio) determine the scaling slope in a dense forest stand. As such, it could provide an independent metric to estimate carbon flux – biomass loss from scaling slope while combining with information from growth (i.e., from remote sensing).

The model (Fig. 1b) predicts that a steeper slope (more negative *α*) should be found in locations with a faster rate of growth relative to biomass loss from mortality, thus leading to a higher rate of living biomass accumulation at the forest stand level. This prediction is applied in forests with even sized structure (i.e., planted forests with no or only minimal intervention after planting) or natural forests with lower environmental heterogeneity or forest patches with even age/size structure) without effects of recruitment. However, the substantial size variation within forest stands could complicate the prediction (23, 24). By linking forest demography with stand structure, the mean tree size vs density ‘scaling law’ integrates forest structure and the functioning of entire forests (6, 21, 25). Understanding this scaling law and how it varies will be crucial for predicting how forest demography and carbon storage respond to environmental gradients or global change through the lens of forest structure. We further hypothesize that variations in the mean size vs density scaling trajectories (slopes) and the temporal biomass accumulation rates across forest biomes vary as a function of forest stand age due to succession and environmental/demographic conditions.

To test these hypotheses, we compiled a large-scale dataset of long-term forest stand observations which were established in largely undisturbed forests covering the period from 1951 to 2019. These forests were distinguished by being in the stage of forest development, i.e., with increasing mean tree size of surviving trees and decreasing total tree density (see Methods and SI Appendix, Methods). These largely undisturbed forests include planted and natural forests in which major disturbances such as fires and human harvesting are not reported during vegetation surveys. Each forest stand had at least four censuses and the stands were located in temperate (n = 800; 92 ha), boreal (n = 602; 64 ha), and tropical (n = 31; 66 ha) forests in North and South America, Europe, and Oceania (SI Appendix, Table S1; SI Appendix, Fig. S1). We quantified the mean tree size vs density scaling trajectory or slope (temporal α) for each forest stand (Fig. 1b), and compared the slopes across sites among and within forest biomes (temperate, boreal and tropical biomes). We investigated the demographic (growth and biomass loss from mortality), forest stand age, and environmental (i.e., climate and soil properties) controls on temporal α in North America, the only region where enough sites were available to cover both the age and rainfall dimensions. We examined the association between temporal α and the biomass accumulation rate across forest biomes. Furthermore, we quantified the spatial α at forest biome scales by using spatial data of mean tree size and density (Fig. 1a). As such, it allowed us to compare the spatial α and mean values of temporal α at forest biome scales.

## Results and discussion

We found considerable variation in the mean tree size vs density scaling slopes (i.e. temporal *α*) across forest stands within each forest biome (−0.2 to -4; Fig. 2). Nevertheless, despite this large variation, we observed a significant difference of the mean value of temporal *α* across forest biomes (both P < 0.001), with less negative slopes in boreal forests (mean ± 1 SE: -1.1 ± 0.03) than in tropical (mean ± 1 SE: - 1.8 ± 0.16) or temperate (mean ± 1 SE: -1.7 ± 0.03) forests. The pattern of temporal α at forest biome scale was robust to the analysis of the bootstrapped (1000 iterations) probability distributions (P < 0.0001) (Fig. 2b). The spread or uncertainty around the mean temporal *α* (mean ± 1 SE: -1.8 ± 0.16) was greatest in tropical forests, where the sample size was lowest (n = 31), thus highlighting a priority to obtain more data in this region. Our approach of estimating the temporal *α* within forest stands differs from the classic and original studies which estimated spatial *α* at macroecological scales (i.e., by using large scale and spatial data of mean tree size – aboveground biomass and density) and predicted the generality of the Euclidean (−3/2) or fractal scaling exponent (−4/3) (4, 14, 15). The variation in temporal *α* in our datasets was comparable to or greater than that found in previous regional-scale analyses which have similarly estimated temporal α within forest stands and suggested a variation of α in the range -0.5 to -3 (22). The considerable variation of temporal *α* across forests of differing age, composition, size structure and resource status suggests that variations in environmental drivers, resource conditions (that might also change with forest development; 26) and forest properties could be thus important in influencing the linkage of mean tree size vs density scaling and functions (27, 28), as explained in detail below.

**Fig. 2.**
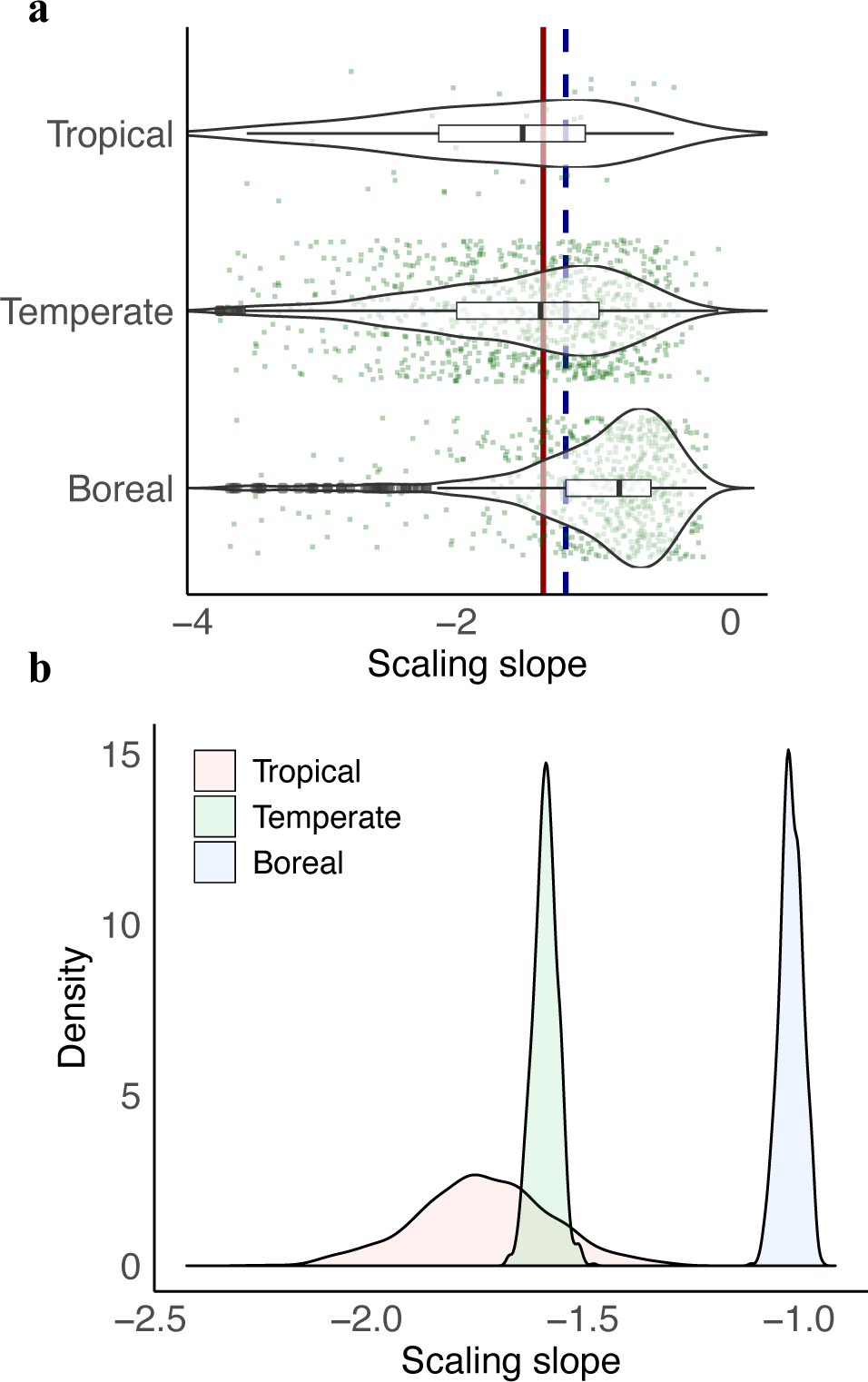
The transient mean tree size vs density scaling exponents (or slopes) over time within and among forest biomes. The scaling slopes across forest stands within tropical (n = 31), temperate (n = 800) and boreal (n = 602) forest biomes (**a**). The solid vertical line refers to the Euclidean scaling law (−3/2) while the dash vertical line refers to the fractal scaling law (−4/3). The bootstrapped (1000 iterations) probability distribution of mean value of the transient mean tree size vs density slope of by randomly selecting 95% of stands across forest biomes (**c**).

To evaluate potential factors influencing the variability in *α*, we focused on North America, which had a large enough sample size for analysis of large vegetation and environmental gradients. Forest stand age and rainfall seasonality were the two most important variables influencing temporal *α* in North American forests. Younger forests and/or forests with lower rainfall seasonality had more negative temporal *α* (Figs. 3a, 3b; SI Appendix, Fig. S3). The two dimensions or interactions of forest stand age and rainfall seasonality largely define the boundary of temporal *α* in North America forest stands, with the most negative temporal *α* values occurring in younger forests with less seasonal precipitation (Fig. 3c). The analysis separating temperate and boreal forests in North America further demonstrated the positive correlations between forest stand age and scaling slopes (Fig. 3d). These patterns were largely driven by the lower tree growth in older forest or forests with higher rainfall seasonality (SI Appendix, Fig. S4), consistent with our conceptual models’ predictions (also see the following section on demographic association with temporal *α*).

**Fig. 3.**
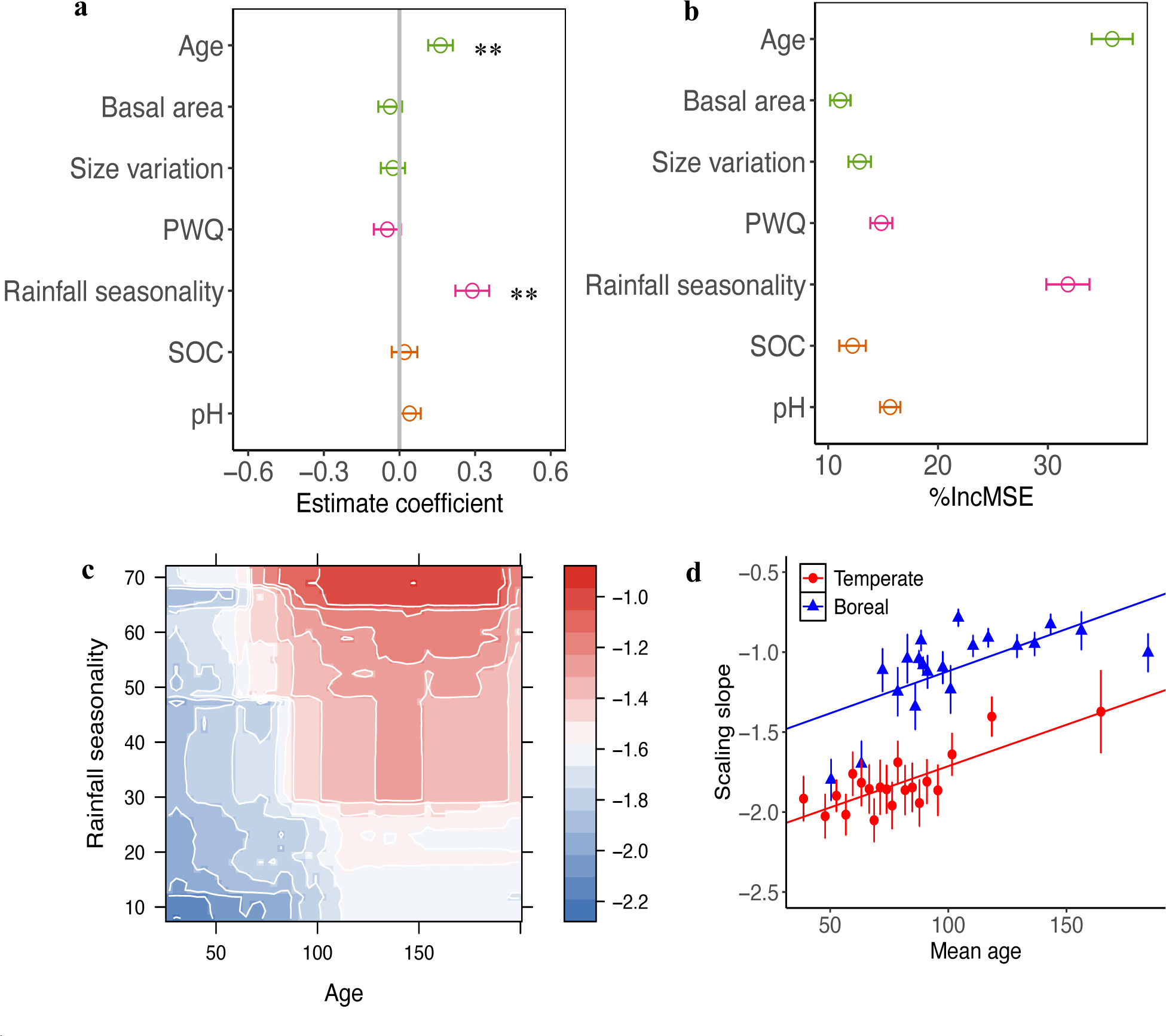
The environmental controls on the transient mean tree size vs density scaling over time in North America forests. **(a)** Standardized coefficient estimates (mean ± 95% CIs) for the effects of forest age, basal area, size variation, precipitation in the warmest quarter (PWQ), rainfall seasonality, soil organic matter (SOC), and pH on the slopes of the transient mean tree size vs density scaling. The environmental variables were standardized (z-score) before analysis. **(b)** Mean decrease in accuracy (%IncMSE, mean and SD) estimated from 1000 simulations of random forests in evaluating the importance of environmental conditions in the slopes of the transient mean tree size vs density scaling. **(c)** Partial feature contributions of primary environmental variable interactions (age vs rainfall seasonality) to the slopes of the transient mean tree size vs density scaling. **(d)** The relationship between forest mean age and the slopes of the transient mean tree size vs density scaling across temperate and boreal forests in North America. Data was binned over 30 points across forest mean age range.

Size inequality, age evenness, and asymmetric competition have been proposed to influence *α* in model simulations and local-scale analyses (23, 24). However, we did not find an influence of tree size variation and other vegetation properties, such as basal area, on α in North America. This could be because at the spatial scale of a landscape or above, including a mosaic of forests across different successional stages with varying forest stand age, species composition and environmental conditions, size structure is either a second order effect or is strongly correlated with the variables describing this mosaic. As such, this further supports the robustness of the prediction of our conceptual model which links temporal *α* with forest demography and carbon dynamics at the ecosystem scale without accounting for tree size structure. Our compiled forest inventory data, which are largely undisturbed, include both planted forests and natural forests in which systematic ecological processes (i.e., asymmetric competition) across forest stands may have masked the role of size variation in a heterogeneous landscape. We do note, however, that theory points to a stronger potential importance of size structure in structurally complex forests (6), such as those found in the tropics, in which we were not able to test with the existing observations.

We further examined the demographic association with *α* across forest biomes. The results using spatial error models (see Methods) showed a consistent negative relationship between forest growth rates (kg m^-2^ yr^-1^) and temporal *α* across forest biomes (Fig. 4a). Biomass loss from tree mortality showed, conversely, a positive relationship with temporal *α*. These results are robust to variations in stand area and data source (i.e., with different minimum tree size threshold and/or tree allometry equations) assessed as random factors using linear mixed models (see Models; SI Appendix, Fig. S5a). Thus, forests with higher growth rates, or lower biomass loss from mortality, or higher growth to biomass loss from mortality ratio (SI Appendix, Fig. S6) have more negative values of temporal *α*, which itself is consistent with a faster rate of stand-level biomass accumulation, thus supporting our model and conceptual framework predictions. Previous studies have found that increases in tree growth accelerate the rate of carbon loss through tree mortality across landscapes, thus leading to uncertainty in the drivers of the forest carbon sink (2, 29, 30). The forest stand data show a general positive relationship between forest growth and biomass loss from mortality across forest biomes (SI Appendix, Fig. S7). However, this relationship is consistently lower than a 1:1 slope, suggesting that increases in forest growth were higher than biomass loss through mortality. This leads to a net increase in forest carbon storage over time, as predicted by our conceptual model.

**Fig. 4.**
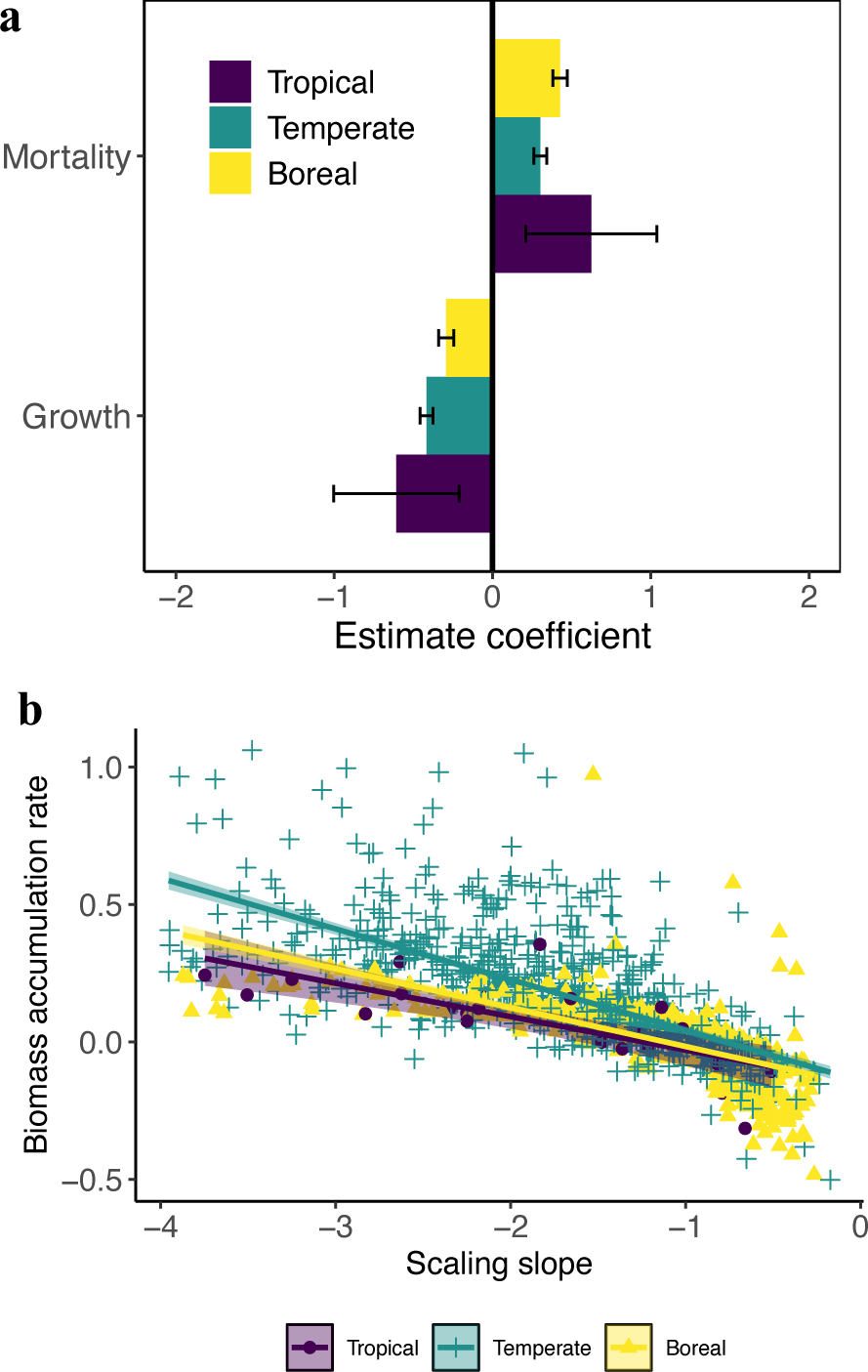
The association of transient mean tree size vs density scaling slopes with the demographic drivers – growth and biomass loss from mortality and biomass accumulation rate across forest biomes. Coefficient estimates (mean ± 95% CIs) for the effects of growth and biomass loss from mortality on the slopes of the transient mean tree size vs density scaling across forest biomes (**a**), quantified by the spatial error models. The relationships between transient mean tree size vs density scaling slopes and biomass accumulation rate (kg m^-2^ y^-1^) across forest biomes (**c**). Both growth and biomass loss from mortality are with units kg m^-2^ yr^-1^. The tails with slope > -1 describe zones of the carbon sources. These results are robust to account for stand area and data source using linear mixed models (see SI Appendix, Fig. S5).

Communities displaying more negative values of temporal *α*, that is, a faster transient trajectory towards fewer and larger trees, also experienced greater rates of living biomass accumulation at a forest stand-scale across forest biomes as shown in both spatial error models accounting for spatial autocorrelation (Fig. 4b) and linear mixed models accounting for stand area and data source as random effects (SI Appendix, Fig. S5b), thus further supporting our model predictions. These relationships between temporal *α* and living biomass accumulation rates at stand level were robustly related to biomass accumulation trends (SI Appendix, Fig. S8). They were also robust to considering stand density and forest stand age (SI Appendix, Figs. S9 and S10). This robustness of pattern was further demonstrated by restricting our analysis to the USA FIA datasets with standardized minimum tree size threshold, forest stand size and tree allometry equations (SI Appendix, Fig. S11), thus highlighting the generality of the relationship between temporal *α* and accumulation rates of living carbon storage across forest biomes.

The ecological consequence of *α* linking forest carbon sequestration and resource availability have remained elusive (4). This variation of the mean tree size vs density scaling slope as a function of stand and environmental parameters has not been shown in previous studies that quantified spatial rather than temporal 𝛼 (4, 15, 25, 31, 32). Indeed, the mean values of temporal 𝛼 and spatial 𝛼 also diverge at forest biome scales (SI Appendix, Fig. S13). The spatial scaling slopes (SI Appendix, Fig. S12) and the temporal transient mean tree size vs density scaling slopes within the same stands are the result of different processes and thus deserve different interpretations (31, 33, 34). The spatial 𝛼 reflects resource availability and functional differences among forest stands (31, 33) implicitly used a space-for-time approach. It, however, does not allow to link the demography with 𝛼 at forest stand or local scales. Instead, the within-stand mean tree size vs density scaling slope or temporal 𝛼 allows to estimate and link both *𝛼* and forest demography for each forest stand and is thus a more powerful and appropriate tool to assess temporal changes in forest functioning (i.e., carbon storage) at forest stand scales.

Nevertheless, the relationship across space between forest growth, biomass loss from mortality and *α* identified here lends support to the hypothesis that *α* may change with changes in resource availability in the future, as induced by environmental changes, resulting in differing forest demography patterns. Increased resource availability associated with global change (including water, increased atmospheric CO_2_ concentration, and increased nutrient availability where from soil organic matter mineralization, biological N fixation or atmospheric deposition) may enhance forest growth and productivity (35, 36). However, in many circumstances global change processes may decrease resource availability (e.g., expected trends in water availability in Mediterranean forests), thus driving declines in forest growth and productivity. Especially, shifts in water availability due to elevated atmospheric demand, changes in precipitation patterns, or increased water use efficiency under elevated atmospheric CO_2_ concentrations (37, 38), may induce shifts in the biomass dynamics of forests. Further quantification of how water use interacts with transient dynamics of mean tree size and density will be required to prove the existence of such resource-driven shifts in dynamics within forest stands (39).

Our results derived from a conceptual framework based on forest demographic processes and long-term forest inventory data, provide linkages between forest structure and forest functions (i.e., demography and biomass accumulation rate) at local or forest stand scales. Our demographic approach reconciles with the previous prediction by the macroecological approach in which growth and mortality rates at broad spatial scale are power functions of tree diameter – the extension of metabolism theory prediction in linking tree allometry with demography (see Methods) (40). However, tree allometry and its linkage with demography vary at local scales in a heterogeneous landscape subjected to environmental and biological influences (18, 19). As such, our demographic approach captures changes in forest structure – mean size and density trajectory derived from forest demography which can be affected by tree allometry and also by local environmental conditions. By linking the changes in individual tree characteristics with local environmental conditions and ecosystem-level dynamics, this refined understanding of mean tree size vs density scaling relationships across different ecosystems can provide mechanistic insights into how carbon dynamics may respond to environmental change.

Our results further highlight the potential to understand forest functions (for carbon storage and dynamics) at broad regional scales by leveraging emerging advanced tools such as high-resolution satellite remote sensing data and size-structured forest demographic models. Advances in high resolution satellite or aerial remote sensing allow surveys of tree-level information (i.e., density, size and allometry) at broad spatial scales (7, 8), although the survey of understories remains challenging in dense and structurally complex natural forests. Another advantage of remote sensing data is its consistent time resolution or scale of vegetation survey across space, thus allowing for the potential of integrating or aggregating data at different spatial scales. Combining forest inventory data with forest stand-to tree-level remote sensing data holds the potential to evaluate forest carbon dynamics (i.e., flux - growth and mortality) from forest structure (i.e., size vs density scaling relations or age) robustly across large scales. Further, the relationships derived here provide both ecological insights and excellent benchmarks (i.e., temporal *α* as a function of forest stand age as shown in Fig. 3d) to facilitate size-structured modelling in land surface models that incorporate demographic processes to assess large-scale biomass dynamics across forests ecosystems under current and future climate change scenarios (9, 11, 41).

Our theoretical framework links forest structure with carbon fluxes - growth relative to biomass loss from mortality in the forest development stage (with increased mean tree size and decreased tree density). When combined with information on tree growth rates (i.e., from remote sensing), the mean tree size vs density scaling relationships quantified here can be used as an independent empirical measure to estimate the expected changes in biomass loss from tree mortality and forest biomass sinks under environmental change at broad spatial scales – the variables which remain highly uncertain in both observations and models. While applying our model and conceptual framework predictions, we suggest caution and more research efforts (theoretically and empirically) in structurally complex forests (especially in tropical forests where appropriate data are currently sparse) in which the role of size structure could interact with major disturbances (i.e., fires and extreme drought) causing substantial mortality during forest development. In such scenarios which are expected to increase with increased disturbance regimes in a changing climate, the estimate of the mean tree size vs density scaling trajectories (slopes) could be subject to increased uncertainty at local scales. As such, efforts would be needed to aggregate data at different spatial and/or temporal scales to allow for the effects of disturbances and size structure to compensate across landscapes. Further, as decomposition influences the time scale of carbon release from dead trees into atmosphere (42), more research on decomposition and the linkage of above and belowground process would support a full cycle assessment of the net carbon sink of forests while linking forest structure and functions.

## Conclusion

Our study builds a theoretical framework with empirical evidence to link forest structure properties – the transient mean tree size and density scaling trajectories (slopes) - with forest demography and biomass accumulation rates. The results demonstrated the strong and negative association between growth and biomass loss from mortality ratios and mean size vs density scaling slopes across forest biomes. Stands with more negative mean size vs density scaling slopes showed higher rates of biomass accumulation, indicative of greater resource availability. Our results highlight the potential for obtaining forest structure properties (i.e., inferred from high resolution remote sensing data or fused into forest demographic models) to improve the prediction of the forest demography, carbon flux and dynamics at broad spatial scales in a changing climate. While applying the theoretical framework, caution would be needed in structurally complex forests, especially in tropical forests where appropriate data are currently sparse.

## METHODS

### Conceptual model linking the transient mean tree size vs density scaling to demographic rates and rate of biomass change

Here we develop a conceptual model to link mean tree size – aboveground biomass vs density scaling trajectory with forest demography and rate of biomass change at ecosystem scales. For any forest, without the effects of recruitment, a relationship between mean individual biomass 𝑀̅ and tree density 𝑁 on the logarithmic scale can be described as:

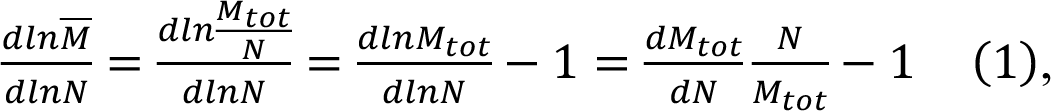

where ***M_tot_*** is total aboveground biomass at ecosystem scale. Similarly, the conceptual model can be also used to link ***M_tot_***vs density scaling trajectory with forest demography and rate of aboveground biomass change at ecosystem scales as, 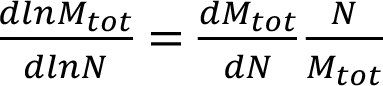. Here we focus on the mean tree size – aboveground biomass vs density scaling trajectory (slope) so that our results could be compared with previous studies (15, 22).

Let 𝜌(𝑚) be the size distribution of a forest, and individual size of this forest ranges from *m_min_* to *m_max_*, we can extend Eq.1 as follow:

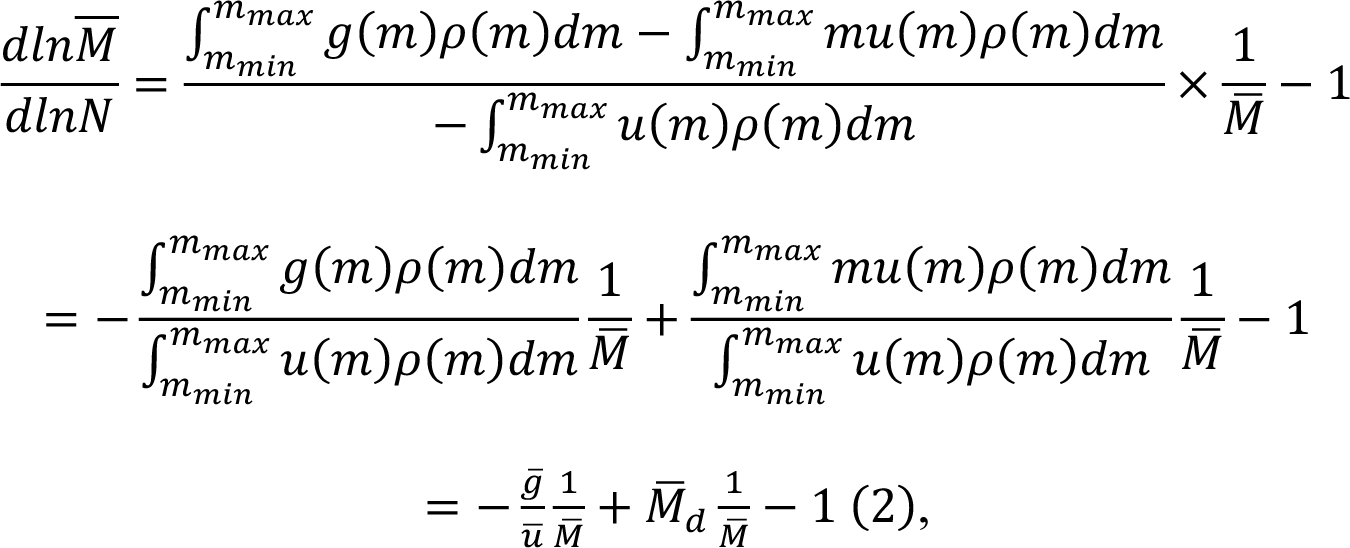

where 𝑔(𝑚) and 𝑢(𝑚) are the growth and mortality rates of an individual of tree size 𝑚, respectively, 𝑀̅, 𝑔̅ and 𝑢̅ are mean alive individual biomass, mean individual growth rate and mean mortality of the forest, respectively. 𝑀̅_*d*_ is mean biomass of dead individuals. This equation thus describes the generalized mean tree size vs density scaling process in which the mean tree size vs density scaling exponent or *α*, regardless of tree size distribution, depends on the averaged forest demography – growth and mortality rates, mean size and mean biomass of dead individuals at community (or ecosystem) scale.

And for 𝑀̅, 𝑔̅, 𝑢̅ and 𝑀̅_*d*_, we have:

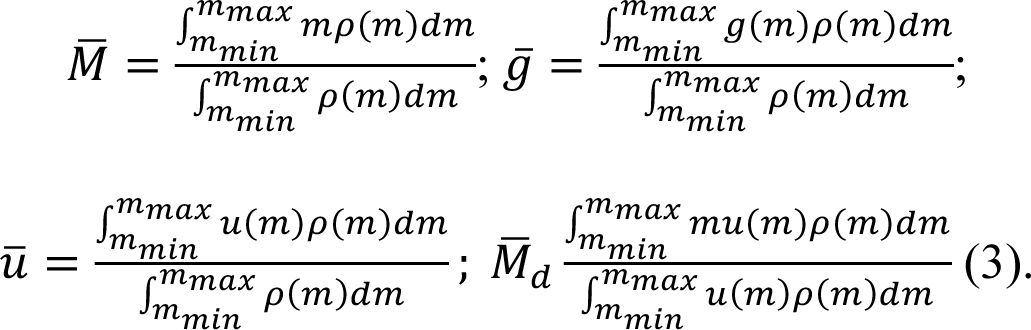

In many classical or original theories of mean tree size vs density scaling, forests (i.e., planted forests or natural forests with lower environmental heterogeneity or forest patches with even age/size structure) are often approximated as composed by equally sized individuals. In such case (even-sized forests), mean tree size vs density scaling relationship or Eq.2 can be described as:

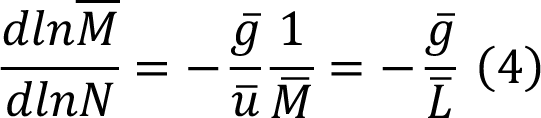

where 𝑔̅ is the averaged growth rate (i.e., kg m^-2^ yr^-1^) and 𝐿̅ is biomass loss (i.e., kg m^-2^ yr^-1^) derived from mortality rate (i.e., ind m^-2^ yr^-1^) multiplying mean size (i.e., kg ind^-1^) at forest stand scale. For even-sized forests, the average growth and mortality rates are also equal to the growth and mortality rates of individuals. At macroecology scales, forest demographic rates have been predicted to be the power functions of size (diameter or mass), i.e., 𝑔̅ = 𝑔_0_𝑀̅^β^ and 𝑢̅ = 𝑢_0_ 𝑀̅^β^, whereby 𝛼 and 𝛽 are scaling exponents as predicted by metabolism (40). As such, our demography approach reconciles with the macroecology approach at broad spatial scales.

Collectively, these eqn derivations suggest that the average growth and biomass loss from mortality jointly or growth and biomass loss from mortality ratio in each forest stand determines the transient mean size vs density scaling trajectory or slope - *α*. When the growth rate is balanced by biomass loss from mortality, the forests could reach the equilibrium state with carbon neutrality (i.e., *α* = -1). A higher ratio, expected for a higher resource condition, could lead to a more negative value of *α* indicating a faster rate of living biomass accumulation rates. The linkage of the mean tree size and density scaling relations with forest demography – growth and mortality motivates us to propose a conceptual framework as demonstrated in Fig. 1b. However, we caution that complications arise in structurally complex natural forests with substantial size variation among individuals (see eqns 2 and 3).

### Forest stand data

The data used in this study were acquired through an extensive literature compilation of long-term forest monitoring sites. The data met the following criteria: 1) The forest stands have at least four censuses and are largely undisturbed by fires and human harvesting, thus allowing for the fitting (through reduced major axis regressions) of the tree mean size vs density scaling trajectory within each forest stand. Tropical forest stands are natural without plantations. By comparison, the largely undisturbed forests in temperate and boreal biomes are either planted or natural forests, but incomplete plantation histories make it challenging to accurately differentiate these stands as planted or natural forests. 2) The tree density averaged over all censuses is large (N > 100 individuals per ha) (15) and stand age (when available) is greater than 25 years. Thus, the forest stand is fully stocked (> 5 kg m^-2^) and is likely to undergo the development or mature stage with decrease in tree density and increase in mean tree size. 3) The density decreases with time and the mean size increases with time (examined through linear regressions). 4) Stands with adequate fits of the mean tree size vs density scaling law (i.e., P < 0.1 and R^2^ > 0.3) were selected.

We clarify that the forest stands selected in this study include those which may have experienced significant changes in climate (i.e. drought or warming) and thus soil resources, and even small local-scale disturbance. Thus, the mean tree size vs density scaling trajectory is jointly governed by competition and environmental (e.g., climate) conditions which allow us to link the scaling slopes with demography and environmental conditions at local or forest stand scales. This is essential because changes in environmental (climate) conditions across space or time, influence the resource footprints and thus the competition among plants. We also clarify that the criteria used above to select the forest stands have substantially reduced the sample size of forest stands because estimate of temporal *α* at each stand scale is subject to challenge and uncertainty deriving from small area size of forest stands, development stages, disturbance, limited number of census and size structure. Table S1 summarizes the number of forest stands compiled in each biome. Further details for the criterion of selected forest stands, forest stand establishment, and measurements were described in SI Appendix, Methods.

### Quantification of the transient mean tree size vs density scaling slopes

Ordinary least squares (OLS) regressions and reduced major axis (RMA) regressions have been widely used in fitting a linear relationship between two variables. Because the mean size and density relationships do not have direct directional causality (43), we used the reduced major axis (RMA) regressions to fit the mean tree size vs density scaling slopes (*α*) at the forest stand scale for log_e_-transformed values of aboveground living vegetation biomass per individual (kg ind^-1^) and tree density (individuals ha^-1^) (44). The fits estimated the slope (*α*), intercept, the significance (P) and the r squared value (R^2^) for each forest stand. The results for each forest stand were then averaged to quantify *α* at scales of forest biomes. To estimate the uncertainty on the means, a bootstrapping approach (1000 iterations), randomly selecting 95% of forest stands, was used to estimate the probability distribution of the mean value of *α* across forest biomes. To account for the low sample size in tropical (South America) forests, the bootstrapped (1000 iterations) probability distribution of mean slope value of *α* was also quantified by randomly selecting 40 across forest biomes.

### Quantification of the demographic rates

In tropical regions, we used the publicly available data of demographic rates – growth and biomass loss from mortality at stand scales. By comparison, in temperate and boreal forests we used the available data (shared by the data providers) such as the individual aboveground woody biomass, tree status (alive, dead, or recruited) to estimate growth and biomass loss from mortality at stand scales. Growth (net woody primary productivity, kg m^-2^ y^-1^) included components of recruitment of new trees and growth of surviving trees and mortality rate (number of individuals per ha per unit time) while biomass loss (kg m^-2^ y^-1^) from mortality was quantified through tree mortality in each census interval.

### The association of transient mean tree size vs density scaling slopes with biomass accumulation rate

To examine whether locations with steeper mean tree size vs density scaling slopes have a higher temporal biomass accumulation rates, we used ordinary least squares regression (OLS) to evaluate the relationships between *α* with biomass accumulation rates, as quantified by the difference of the growth (kg m^-2^ y^-1^) and biomass loss (kg m^-2^ y^-1^) averaged over censuses. To test the sensitivity, we also used OLS to fit biomass with time (year) of each census and quantify biomass temporal trends (SI Appendix, Fig. S2). The forest stands with adequate OLS fittings (i.e., P < 0.1 and R^2^ > 0.3) were selected to examine these relationships between *α* and biomass temporal trends. To account for the potential effects of density (see Eqn 1 in main text), we also included 𝑁*^α^* or 𝑁 in the OLS models to examine the relationships between *α* and temporal biomass accumulation rates. To account for the potential effects of small plot size (i.e., < 0.1 ha) and varying data source with different criteria of different minimum tree size threshold and/or tree allometry equations across regions, we also used linear mixed model to account for stand area and data source as random effects.

### Predictors of the transient mean tree size vs density scaling slopes

We examined the influence of environmental conditions (vegetation, climate, and soil properties) (see SI Appendix, Methods) and forest demographics, the growth (kg m^-2^ y^-1^) and biomass loss from mortality (kg m^-2^ y^-1^) as well as growth and biomass loss from mortality ratio (see Eqns 2 and 4), on the mean tree size vs density scaling slopes. We focused this analysis in North America where we had a large sample size with large environmental gradients and more importantly with available forest stand age information. We clarified that we averaged the forest growth and biomass loss from mortality over the whole censuses in each forest stand, consistent with the quantification of the mean tree size vs density scaling slopes. Thus we investigated the association of overall growth and biomass loss from mortality and growth and mortality ratio averaged over the time window of the trajectory in mean tree size vs density scaling slopes and biomass accumulation rates across landscapes. To account for spatial autocorrelation, spatial error models (SEM) (45) were used to examine the environmental and demographic drivers of mean tree size vs density scaling slopes, respectively (SI Appendix, Methods). Moreover, linear mixed model was used to account for stand area and data source as random effects while investigating the demographic drivers of mean tree size vs density scaling slopes. Furthermore, a random forest machine-learning algorithm was used to determine variable importance for environmental drivers (46) (SI Appendix, Methods). Growth was log-transformed to meet the assumptions of normality of residuals when examining its role in mean tree size vs density scaling slopes. Diagnostic analyses of homoscedasticity, multivariate normality and independence of residuals were also conducted to test the assumptions of the models (47).

## Data and code availability

Data of mean size vs density scaling trajectories (slopes) and the underlying environmental and biological variables in each forest stand as well as codes will be deposited in github and publicly available deposit such as Dryad when the study is accepted. The raw inventory data are available upon reasonable request from the corresponding author.

## Author contributions

K.L.Y. and T.A.M.P. designed research; H.Y.H.C., A.G., E.B.S., R.B.A., H.P., P.C., O.L.P., R.J.W.B., C.J.C., S.B.X and A.P.B. contributed new data and analytic tools; K.L.Y. analyzed data; K.L.Y., T.A.M.P., and A.P.B. wrote the paper with inputs from all coauthors.

## Acknowledgements

Acknowledgemeants

T.A.M.P. acknowledges funding from the European Research Council (ERC) under the European Union’s Horizon 2020 research and innovation programme (grant agreement no. 758873, TreeMort). This study is a contribution to the strategic research areas BECC and MERGE and the Nature-based Future Solutions profile area.

