## SUPPORTING ONLINE MATERIAL for "Forest demography and biomass accumulation rates are associated with transient mean tree size vs density scaling relations"

### Forest stand data

The majority of stands which meet the criterion of quantifying transient mean tree size vs density scaling exponents were from Canada and the Forest Inventory and Analysis (FIA) Program of the U.S. Forest Service. The stands from Canada were located in Alberta (AB), Saskatchewan (SK), Manitoba (MB), Ontario (ON) and Quebec (QC). They were compiled based on previous studies (1–3) and we here provided a summary for these stands. The stands were established in visually homogenous well-stocked stands that were at least 1 ha in size. The stands were located at least 100 m from any openings to minimize the impacts of edge effects. Each stand was of a fixed area (area dependent on jurisdiction) and all trees within the stand that met the diameter threshold (threshold dependent on jurisdiction) were tagged and had their species identification recorded. Stands where trees were inaccurately recorded (i.e., trees were re-numbered each census) were excluded from the dataset. Canada forest stands have good records of vegetation regeneration history. Only the stands with natural regeneration were selected and thus the stands were largely unmanaged. Some forest stands in QC have low biomass (i.e., <5 kg $m^{-2}$) and were thus excluded in the analysis. For the purpose of our study, we used the available data such as the aboveground living vegetation (AGB) (kg $m^{-2}$), living vegetation density (N, number of individuals per ha), forest age, basal area, and species richness at each census. We then used reduced major axis (RMA) regressions (4) and ordinary least squares regression (OLS) to fit the transient mean tree size vs density scaling over time. With the information of tree recruitment and tree death being available (see Supplementary Table2), we also quantified the forest growth (net woody primary productivity, kg $m^{-2}$ $y^{-1}$) including components of recruitment of new trees and growth of surviving trees and mortality rate (number of individuals per ha per unit time) while biomass loss (kg $m^{-2}$ $y^{-1}$) was quantified through tree mortality in each census interval, following the methods reported in the previous studies (1–3).

The rest majority of forest data were from the Forest Inventory and Analysis (FIA) Program of the U.S. Forest Service. With its open access, FIA data has been widely used in previous studies (5, 6). Thus, here we provide a summary for these stands. Applying a nationally standardized sampling protocol, FIA inventory stands in forested areas consist of four 7.2 m fixed-radius substands spaced 36.6 m apart in a triangular arrangement with one substand in the center. FIA stands had records in each census of the diameter of breast height (DBH) of all individual trees above a defined diameter (12.7 cm) at breast height threshold. The threshold of 12.7 cm is higher than those in Canada stands and thus may underestimate the growth because of limited records of recruitments. We extracted the full FIA dataset with at least four censuses and excluded stands that reported any human-caused disturbances, such as fire, logging. As compared to Canada data, the record of stand regeneration history (i.e., logging) is not as good as FIA forest stands. Thus, to further make sure that the FIA stands used in the study are largely unmanaged, we used the high resolution Global 1-km Consensus Land Cover map (7), which reported the vegetation types including cultivated and managed vegetation. The stands with cultivated and managed vegetation cover more than 30% were thus excluded in this study. In total, we compiled 490 stands located in eastern United States ranging from 1997 to 2019, which met the criteria of our study (see main text).

The majority of forest stand data in Europe were from the International Co-operative Programme on Assessment and Monitoring of Air Pollution Effects on Forests (ICP Forests) launched in 1985 under the UNECE Convention on Long-range Transboundary Air Pollution (CLRTAP). The standard protocol for vegetation survey and tree measurements is described in Dobbertin and Neumann (2016). The data which met our criterion of quantifying the transient mean tree size vs density scaling exponent included 61 stands located in Italy, Switzerland, Germany, Spain, Austria, Slovak, France, and Denmark. The rest of forest stand data in Europe was compiled from the previous study (8), which included 21 stands located in Germany meeting the criteria of our study (see main text). Similarly with analysis in Canada and FIA forest stands, we quantified the forest growth (net primary productivity-NPP, kg $m^{-2}$ $y^{-1}$), mortality rate (number of individuals per ha per unit time), and biomass loss (kg $m^{-2}$ $y^{-1}$) from mortality. The available aboveground living vegetation biomass (AGB) (kg $m^{-2}$) and living vegetation density (N, number of individuals per ha) were used to quantify the slope of transient mean tree size vs density scaling.

The stands from Oceania were located in New Zealand's South Island. These stands have been used to study compositional and structural changes in a montane forest at decadal time scales. The 155 stands selected for this study sample this forest along randomly located transects. Each plot was 0.04 ha and used a standard inventory method for New Zealand's natural forests. All tree stems with a diameter at breast height (1.4 m) ≥30 mm were tagged and had their diameter recorded by species in 1974. Recruitment, growth, and mortality were determined using eight subsequent re-measurements of tagged individuals by 2009. The stands were located in forest that has not previously been cleared, burned, of logged.

The tropical forest data were from the previous study (9), which included mature forests across the lowland tropical areas of South America. The data used allometric equations including woody density, diameter and tree height (10) to convert DBH into biomass. It also has the records of tree recruitment and tree mortality, thus allowing us to examine the role of demographic rates (NPP, mortality rate and biomass loss) in the transient mean tree size vs density scaling slopes and forest biomass accumulation rates. For the purpose of this study, we used aboveground living vegetation biomass (AGB) at the start of the census (kg $m^{-2}$), annual net change in AGB (kg $m^{-2}$ $y^{-1}$), and interval time (y) between consecutive censuses to derive the biomass in each census. Total annual AGB mortality (plus added unobserved components) (kg $m^{-2}$ $y^{-1}$), total annual AGB productivity of surviving trees plus recruitment (plus added unobserved components) (kg $m^{-2}$ $y^{-1}$), and interval time between consecutive censuses were used to determine NPP (kg $m^{-2}$ $y^{-1}$) and biomass loss from mortality (kg $m^{-2}$ $y^{-1}$) in each census interval, respectively. For more details, the readers can refer to the study by Brienen et al. 2015 (9), which used this data to quantify temporal trends of tree mortality rates, biomass loss from mortality and forest growth (NPP).

**Predictors of transient mean tree size vs density scaling slopes**

This section provides the details of examining how transient mean tree size vs density scaling slopes were affected by vegetation, climate and soil conditions. We focused on the analysis in North America which has a large sample size and thus spans a large environmental gradient. With respect to forest vegetation properties, we calculated the coefficient of variation (CV) of diameter as the measure of size variation at each forest stand, basal area density (BA) and species richness (Diversity; the number of species) from forest stand data, which have been found to largely affect plant competition (11, 12) and thus may influence the transient mean tree size vs density scaling slopes. BA was normalized by dividing stand area. CV, BA and Diversity across all censuses were averaged to be representative of the averaged conditions of vegetation properties, consistent with the estimate of climate and soil data in each forest stand. Data of forest age was available in Canada and FIA forest stands and this further allowed us to evaluate the effects of forest age in the transient mean tree size vs density scaling slopes. We clarified that the age at each stand is determined by coring dominant or co-dominant trees that represent a plurality of non-overtopped trees. The stand age is thus estimated as the average of the dominant trees (5), assuming that the age of the dominant or codominant trees represents the age of the forest ecosystem.

Based on previous studies (13–16), climate variables which may affect the transient mean tree size vs density scaling slopes were selected for this study. They included annual mean temperature (MAT), annual total precipitation (MAP), precipitation in driest quarter (PDQ), precipitation in warmest quarter (PWQ), and rainfall seasonality (RS, expressed as a coefficient of variation of rainfall). MAT, MAP, PDQ, PWQ, and RS at each forest stand scale were derived from WorldClim variables with a spatial resolution of 1km based on forest stands' georeferenced locations (latitude and longitude). The global rasters of soil organic carbon (SOC), pH and cation exchange capacity (CEC) at 2 m soil depth with 1 km spatial resolution, which play important roles in nutrient cycling (Schmidt et al. 2011), were used to derive soil properties in each forest stand; data of SOC, pH and CEC were downloaded from https://www.soilgrids.org.

The ordinary least squares regression (OLS) models were used to examine which vegetation (age, size variation - CV, basal area, and diversity), climate (MAT, MAP, PDQ, PWQ, and RS), and soil (SOC, pH and CEC) variables were important in affecting the transient mean tree size vs density scaling slopes across forest stands. Prior to data analysis, these variables were examined to avoid multicollinearity using a matrix of pairwise correlations to remove any variable with high correlations ($R>0.7$) with other predictor variables (17). The variables (age, basal area, CV, PWQ, RS, SOC, and pH) that gave the best prediction of variations of the transient mean tree size vs density scaling slopes across space were retained in the model. All of the environmental variables were standardized (z-score) before analysis. The OLS models showed considerable spatial autocorrelation in the residuals in North America forests (Moran's I test: I = 0.13, P < 0.001). We thus used the spatial error models (SEM) to remove the impacts of the spatial autocorrelation in the residuals (Moran's I test: I = -0.023, P = 0.87). To this end, we used a spatial weights matrix with neighbourhoods defined as cells within certain distance of the focal cell. The distance was determined in each forest biome so that 95% of the stands had at least one neighbourhood (18). Some of the forest stands had the same coordinates and we thus generated normal random noise (mean = 0; variance = 0.0001) into the coordinates for quantifying the spatial weights matrix. The similar spatial error models (SEM) were also used to examine the role of demographic drivers in the transient mean tree size vs density scaling slopes (see main text).

To further examine the variable importance of environmental conditions, we used a random forest machine-learning algorithm (19). We ran 1000 simulations of machine-learning algorithm random forest and reported mean values of mean decrease in accuracy (%IncMSE) with 95% confidence interval. The greater the values of %IncMSE are, the more important the variables are.

**Supplementary Table1 Summary of the compiled long-term forest monitoring stand dataset ranging from 1951 to 2019 over at least four censuses across forest biomes.**

| Biomes/ Continents | Number of stands | Total area (Ha) | Earliest census (y) | Latest census (y) | Data source and/or provider |
|---|---|---|---|---|---|
| Tropical/South America | 31 | 66 | 1979 | 2012 | Brienen et al (2015); Pillet et al (2018); |
| Temperate | 800 | 92 | 1951 | 2019 | FIA Forests; ICP Forests (cf., Dobbertin and Neumann (2016)); Yu et al (2019); Rob Allen, Hans Pretzsch, |
| Boreal | 602 | 64 | 1958 | 2015 | Alberta Ministry of Agriculture - Forestry and Rural Economic; Development Saskatchewan Ministry of Environment; Manitoba Department of Agriculture and Resource Development; |

Note: Only the data derived from Pillet et al (2018) do not have forest growth, tree mortality and biomass loss from mortality.

**Supplementary Table2 Summary of quantifying the slopes and intercepts of the transient mean tree size vs density scaling using reduced major axis regressions across forest biomes.**

| Biome/Continents | Slope | Std_error | Intercept | Std_error | Num_stand |
|---|---|---|---|---|---|
| Tropical/South America | -1.8 | 0.16 | 17.7 | 0.99 | 31 |
| Temperate | -1.7 | 0.03 | 16.1 | 0.18 | 800 |
| Boreal | -1.1 | 0.03 | 12.7 | 0.18 | 602 |

Note: Std_error, standard error; Num_stand, number of stands

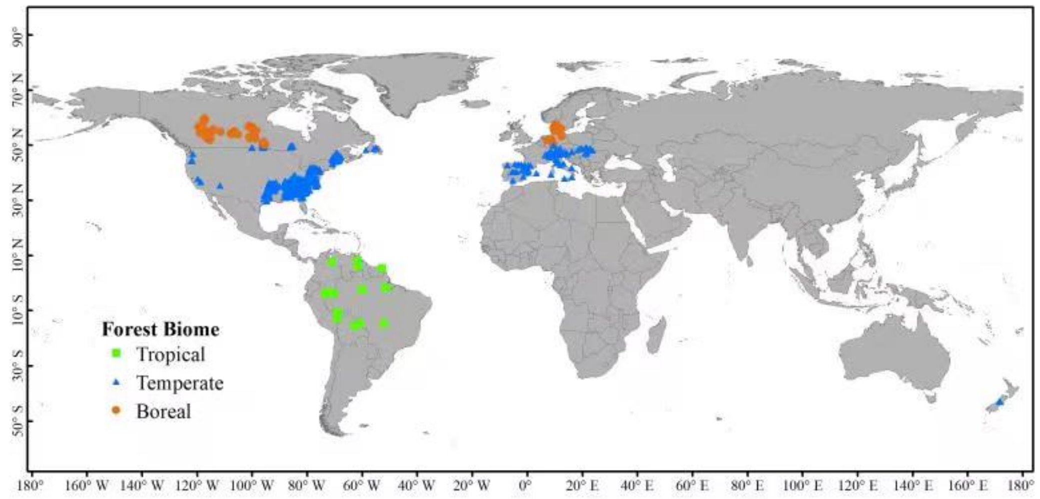


**Supplementary Fig. 1** Long-term forest stand data ranging from 1951 to 2019 over at least 4 censuses across tropical (n = 31), temperate (n = 800), and boreal forests (n = 602).

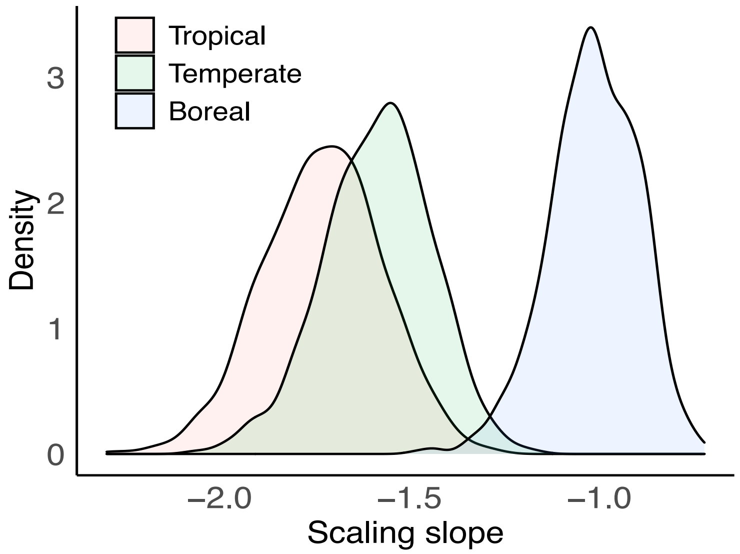


**Supplementary Fig. 2** The bootstrapped (1000 iterations) probability distribution of mean value of transient mean tree size vs density scaling slopes by randomly selecting 30 stands across forest biomes. This is used to account for effects of the low sample size in tropical (South America) forests.

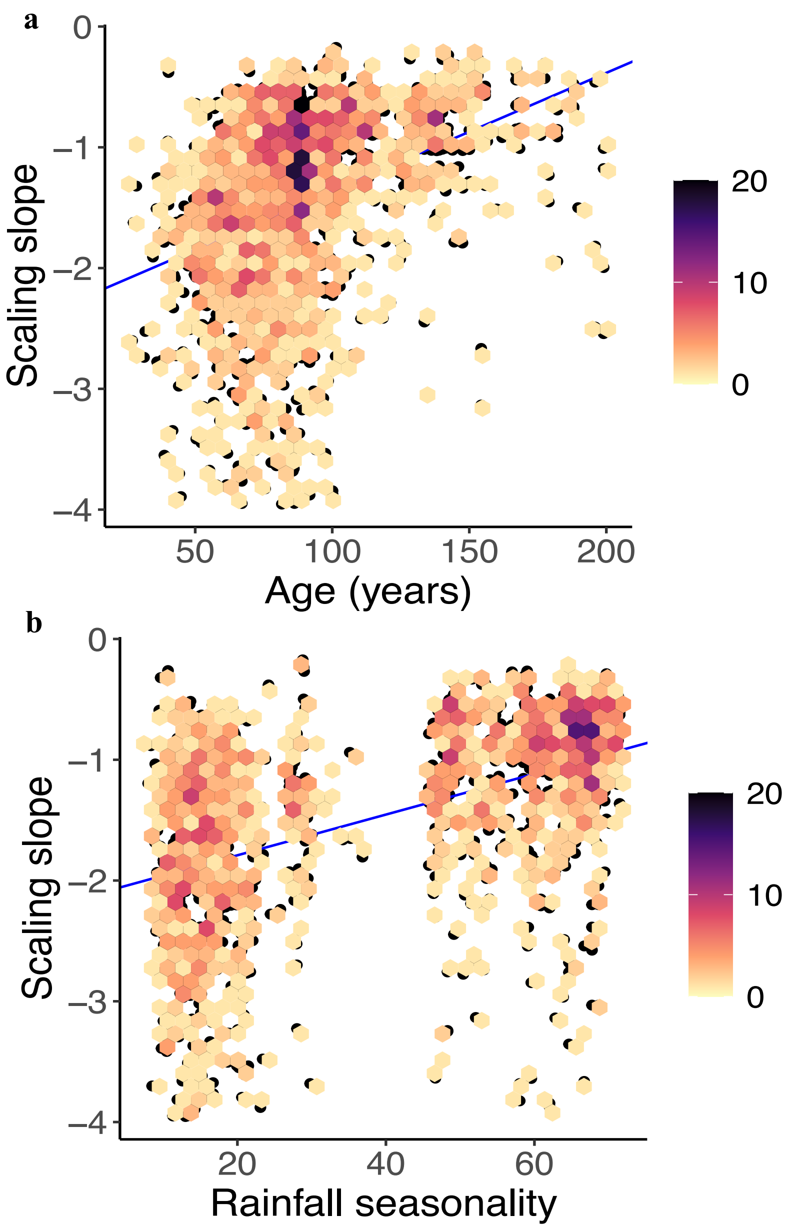


**Supplementary Fig. 3** The relationship between slopes of transient mean tree size vs density and forest age (**a**) and rainfall seasonality (**b**) in North America forests.

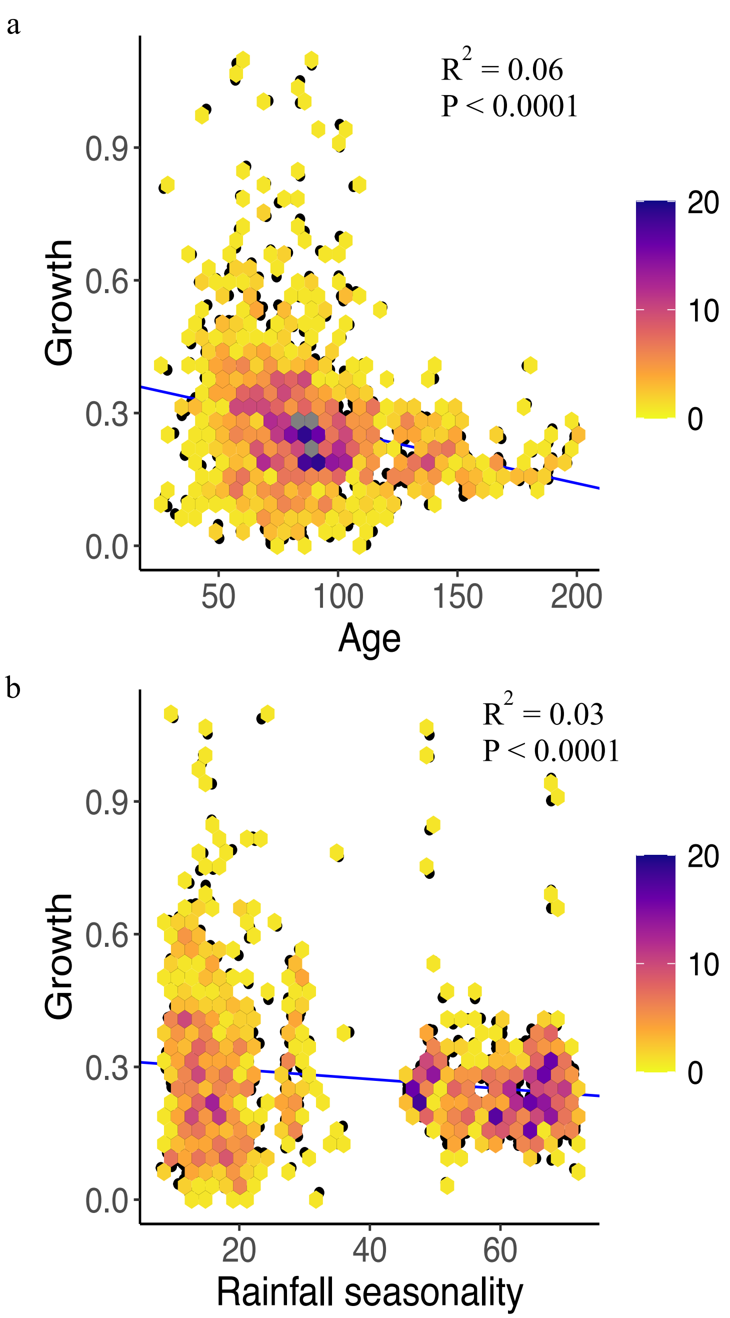


**Supplementary Fig. 4** The relationship between growth (kg $m^{-2}$ $yr^{-1}$) and forest age (**a**, **years**) and rainfall seasonality (**b**) in North America forests.

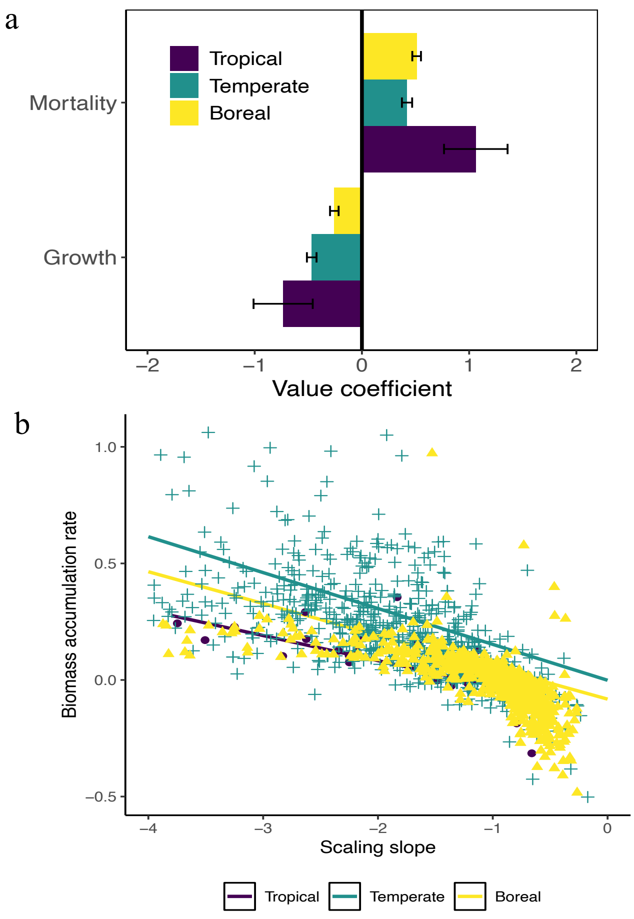


**Supplementary Fig. 5 The association of transient mean tree size vs density scaling slopes with the demographic drivers – growth and biomass loss from mortality and biomass accumulation rate across forest biomes.** Coefficient estimates (mean ± 95% CIs) for the effects of growth and biomass loss from mortality on the slopes of the transient mean tree size vs density scaling across forest biomes (**a**), quantified by linear mixed models which account for stand area and data source (i.e., with different minimum tree size threshold and/or tree allometry equations) as random factors. The relationships between transient mean tree size vs density scaling slopes and biomass accumulation rate (kg $m^{-2}$ $y^{-1}$) across forest biomes (**c**). Both growth and biomass loss from mortality are with units kg $m^{-2}$ $yr^{-1}$. The tails with slope > -1 describe zones of the carbon sources.

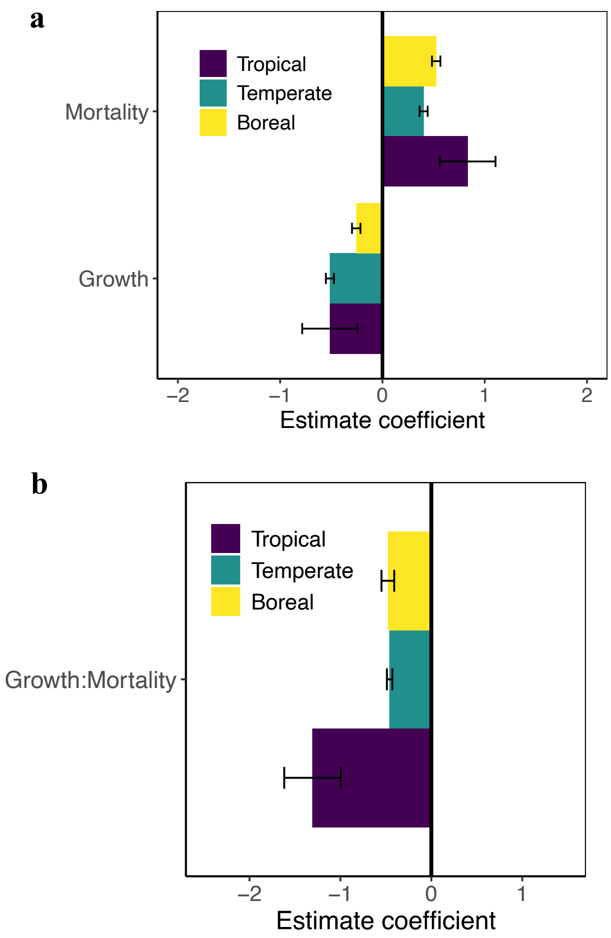


**Supplementary Fig. 6** Standardized coefficient estimates (mean ± 95% CIs) for the effects of growth and mortality, or growth and mortality ratio on the slopes of the transient mean tree size vs density scaling across forest biomes (**a, b**), quantified by the spatial error models. Both growth and mortality are with unit of units of kg $m^{-2}$ $y^{-1}$ and were standardized (z-score) before analysis.

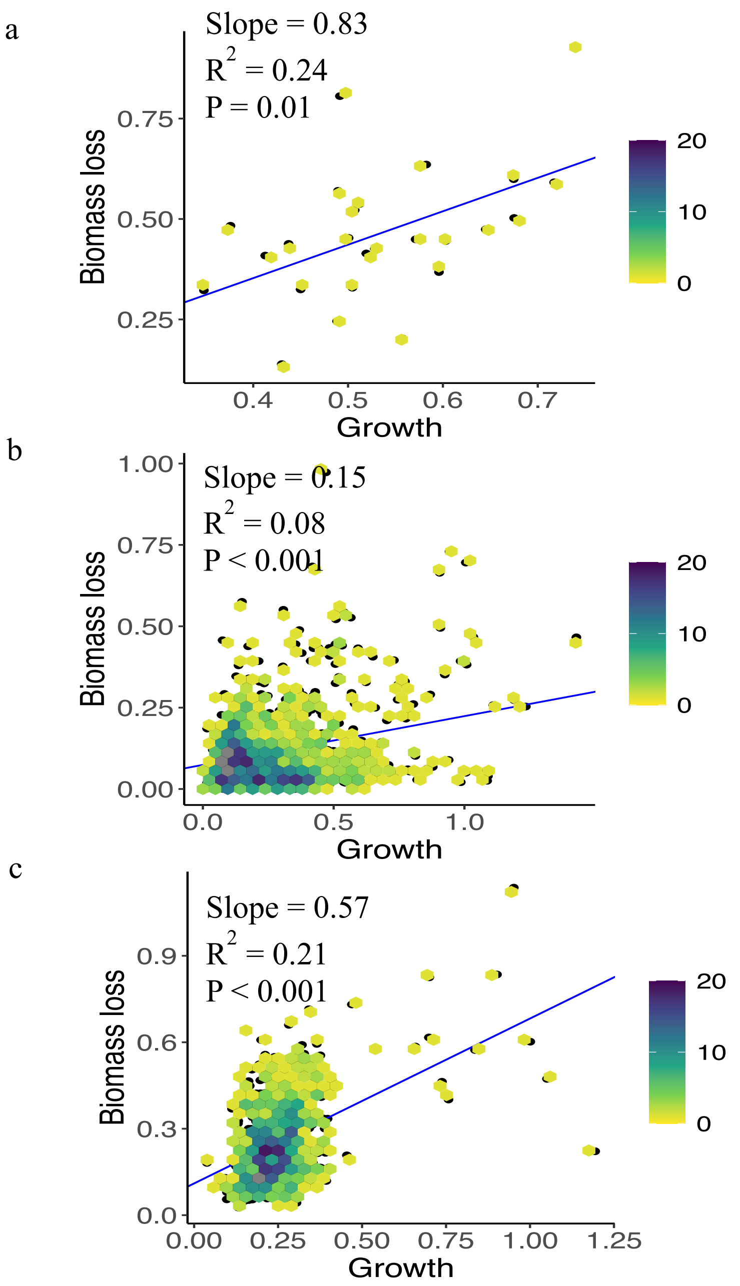


**Supplementary Fig. 7** The relationships between forest growth (kg $m^{-2}$ $y^{-1}$) and biomass loss (kg $m^{-2}$ $y^{-1}$) from mortality across tropical (**a**), temperate (**b**) and boreal (**c**) forests. Forest growth and biomass loss were averaged over the whole inventory period to be representative of the averaged conditions of vegetation dynamics, corresponding with the quantification of the transient mean tree size vs density scaling over time.

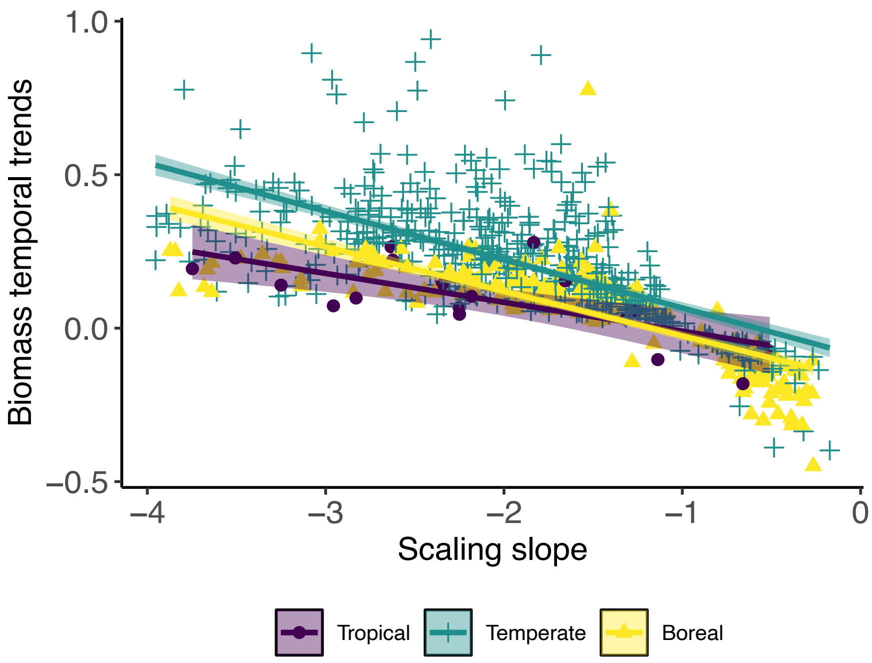


**Supplementary Fig. 8** The relationships between the slopes of the transient mean tree size vs density scaling and temporal trends of biomass (kg $m^{-2}$ $y^{-1}$) across forest biomes. Temporal trends of biomass were estimated by using ordinary least squares regression (OLS) to fit biomass with time (year) of each census (see Methods).

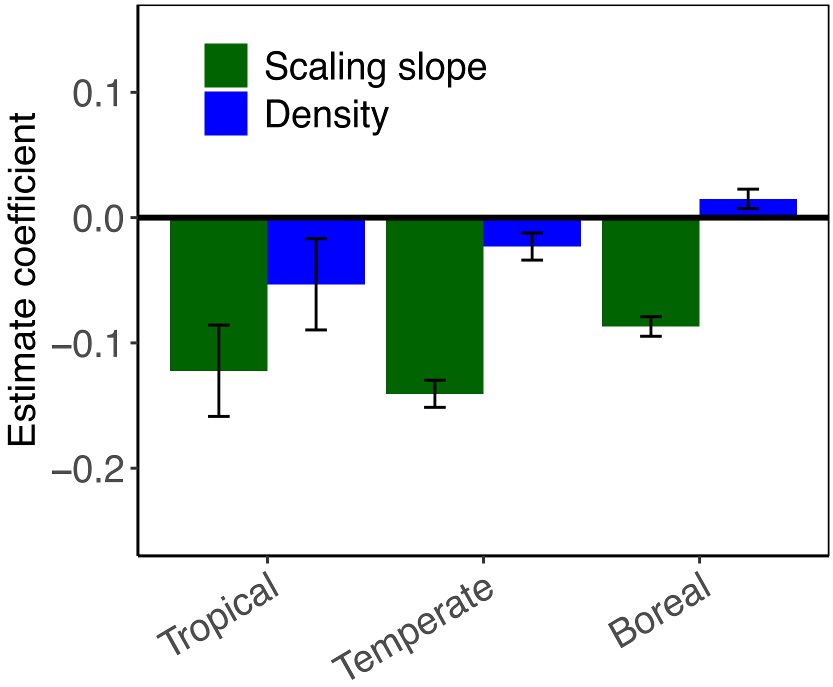


**Supplementary Fig. 9** The role of transient mean tree size vs density scaling slopes in biomass accumulation rates by accounting for the effects of tree density across forest biomes. Coefficient estimates (mean ± 95% CIs) of the relationships between the slopes of the transient mean tree size vs density scaling and density ($N^{\alpha}$) and biomass accumulation rates (kg m$^{-2}$ y$^{-1}$), quantified as the difference of the growth (kg m$^{-2}$ y$^{-1}$) and biomass loss (kg m$^{-2}$ y$^{-1}$) averaged over censuses across forest biomes. Note that $N^{\alpha}$ was incorporated into the linear regression fittings between transient mean tree size vs density scaling slopes and biomass accumulation rates to account for the potential non-linear effects of tree density. The results were robust to the case of incorporating $N$ into the linear regression fittings.

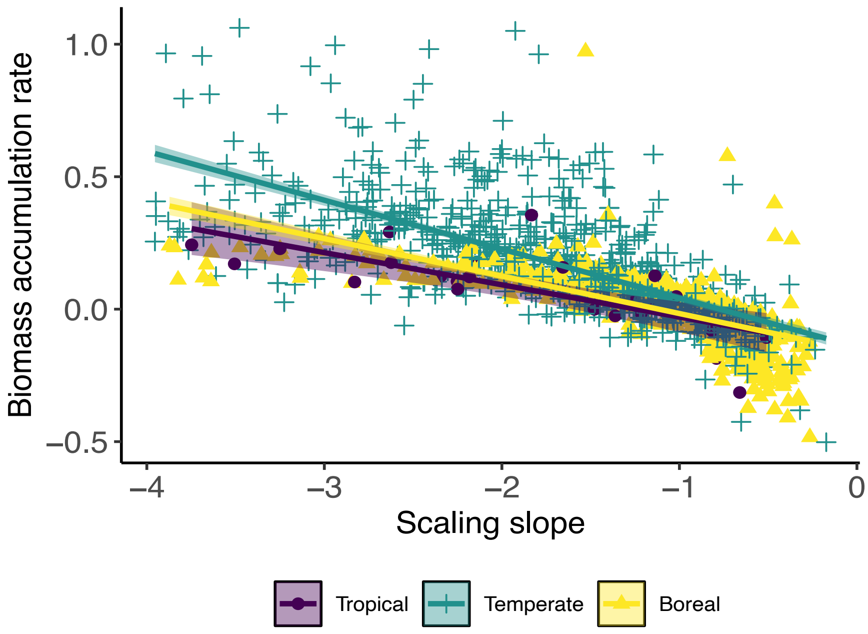


**Supplementary Fig. 10** The relationships between the slopes of the transient mean tree size vs density scaling and biomass accumulate rates (kg $m^{-2}$ $y^{-1}$), quantified as the difference of the growth (kg $m^{-2}$ $y^{-1}$) and biomass loss (kg $m^{-2}$ $y^{-1}$) averaged over censuses across forest biomes, for old growth forests (age >100 years) across forest biomes.

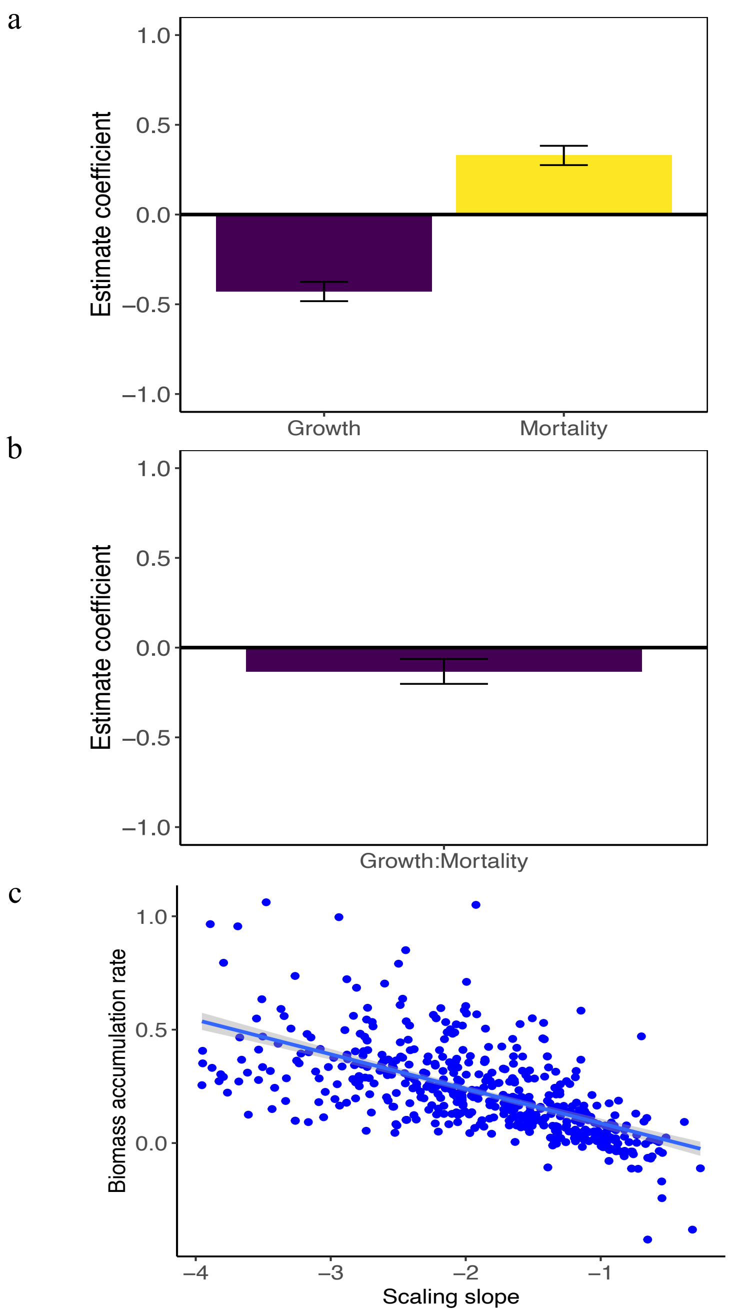


**Supplementary Fig. 11** The strong association of mean tree size vs density scaling slopes with forest demography (a, b) and biomass accumulation rate (c) in USA using FIA dataset with standardized minimum tree size threshold, stand size and tree allometry equations. a and b were fitted by spatial error models with reporting estimate coefficient while c was fitted by ordinary least squares regression.

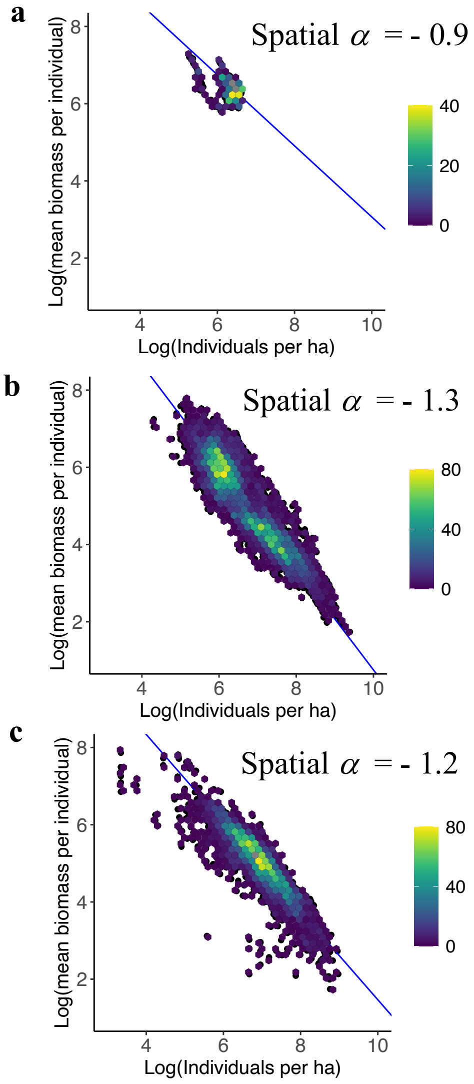


**Supplementary Fig. 12** The interspecific (spatial) scaling relationships between tree density ($\log_e$-transformed) and mean tree size ($\log_e$-transformed) in tropical (**a**), temperate (**b**) and boreal (**c**) forests, quantified by aggregating the data of mean tree size and density across space and time.

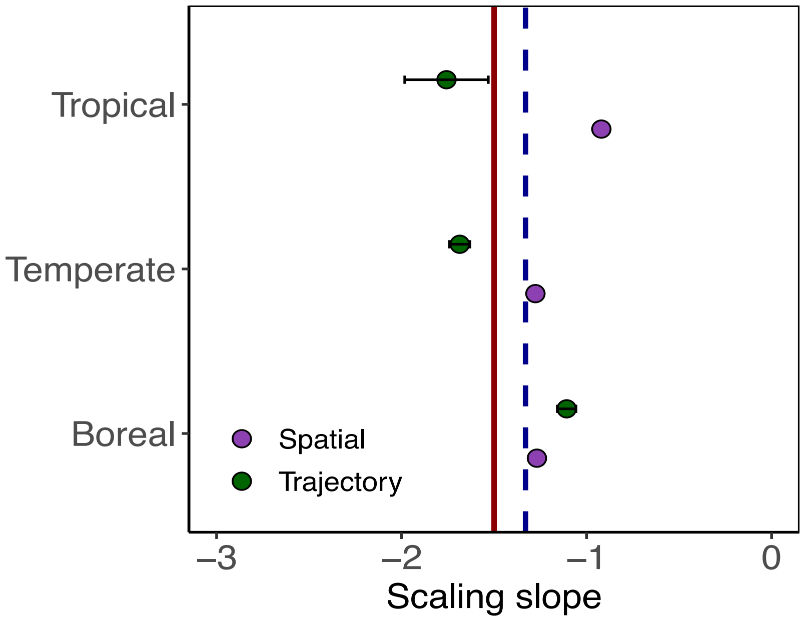


**Supplementary Fig. 13** The slopes of the transient mean tree size vs density scaling over time and spatial scaling slopes across forest biomes. The slopes of the transient mean tree size vs density scaling are mean ± 95% CIs. The spatial scaling slope is one single value in each forest biome or each continent, quantified by aggregating the data of tree density ($\log_e$-transformed) and mean tree size ($\log_e$-transformed) across space and time.
